# Streptolysin-induced endoplasmic reticulum stress promotes group A streptococcal *in vivo* biofilm formation and necrotizing fasciitis

**DOI:** 10.1101/183012

**Authors:** Anuradha Vajjala, Debabrata Biswas, Kelvin Kian Long Chong, Wei Hong Tay, Emanuel Hanski, Kimberly A Kline

## Abstract

Group A *Streptococcus* (GAS) is a human pathogen that causes infections ranging from mild to fulminant and life-threatening. Biofilms have been implicated in acute GAS soft-tissue infections such as necrotizing fasciitis (NF). However, most *in vitro* models used to study GAS biofilms have been designed to mimic chronic infections and insufficiently recapitulate *in vivo* conditions and the host-pathogen interactions that might influence biofilm formation. Here we establish and characterize an *in vitro* model of GAS biofilm development on mammalian cells that simulates microcolony formation observed in a murine model of human NF. We show that on mammalian cells, GAS forms dense aggregates that display hallmark biofilm characteristics including a three-dimensional architecture and enhanced tolerance to antibiotics. In contrast to abiotic-grown biofilms, host-associated biofilms require the expression of secreted GAS streptolysins O and S (SLO, SLS) resulting in the release of a host-associated biofilm promoting-factor(s). Supernatants from GAS-infected mammalian cells or from cells treated with endoplasmic reticulum (ER) stressors restore biofilm formation to an SLO and SLS null mutant that is otherwise attenuated in biofilm formation on cells, together suggesting a role for streptolysin-induced ER stress in this process. In an i*n vivo* mouse model, the streptolysin-null mutant is attenuated in both microcolony formation and bacterial spread, but pre-treatment of softtissue with an ER-stressor restores the ability of the mutant to form wild type like microcolonies that disseminate throughout the soft tissue. Taken together, we have identified a new role of streptolysin-driven ER stress in GAS biofilm formation and NF disease progression.

**Significance Statement:** Although it is well-accepted that bacterial biofilms are associated with many chronic infections, little is known about the mechanisms by which group A *Streptococcus* (GAS) biofilms contribute to acute soft tissue-invasive diseases like necrotizing fasciitis (NF). In this study, we establish a physiologically relevant *in vitro* model to study GAS biofilm formation on mammalian cells and validate our findings in a mouse model that mimics human NF. This study demonstrates a novel role of GAS streptolysin-mediated ER stress in the development and spread of GAS biofilms in acute softtissue infections. We also show that biofilm formation depends on the release of a host-associated factor that promotes microcolony formation and GAS dissemination *in vivo*.

## Introduction

Group A *Streptococcus* (GAS) causes infections ranging from superficial and self-limiting, such as impetigo and pharyngitis, to life-threatening and highly invasive diseases such as streptococcal toxic shock syndrome (STSS) and necrotizing fasciitis (NF) (1, 2). In NF, the infection usually starts as a superficial skin lesion and is rapidly followed by massive bacterial accumulation in the form of dense aggregates that spread along the fascial planes (3). The dissemination of GAS into soft tissue causes severe damage that often results in the mortality of the patient despite prompt medical intervention (3).

Biofilm formation is one of many pathogenic strategies that bacteria employ for disease progression and persistence in hosts (4). Bacterial biofilms are densely packed microbial communities, encased in a self-produced matrix, that are often recalcitrant to antibiotic or immune clearance (5). The presence of GAS communities embedded in a glycocalyx in a murine model of impetigo (6) and in a zebrafish model of myositis (7) suggests that GAS forms tissue-associated microcolonies during the course of infection. Furthermore, three-dimensional GAS biofilm-like structures were observed in a chinchilla middle ear model of human otitis media by scanning electron microscopy (SEM) (8). Antibiotic treatment failure against susceptible clinical GAS isolates has been attributed to biofilm formation, given the increased tolerance of biofilm bacteria to antibiotics (9, 10). Indeed, dense clusters of GAS microcolonies have been observed within the soft tissue of NF patients (11) and GAS isolated from human necrotizing soft tissue infections (NSTIs) appeared as multi-layered fibrous biofilms that persisted despite prolonged antibiotic therapy (12). However, despite evidence of biofilm-like communities during GAS infection, most research on the contribution of biofilms to the pathogenesis of GAS disease has been performed *in vitro,* where infection-associated environments are not fully recapitulated (13).

In the current study, we developed an *in vitro* model of GAS biofilm formation on mammalian cells to study acute, biofilm-associated infection. We demonstrate that, when cultured on mammalian cells but not on abiotic surfaces, the expression of streptolysin O or S (SLO or SLS) is essential for the production of host-associated GAS biofilms. Previously, it was shown that secretion of SLO and SLS led to the release of asparagine (ASN) from host cells due to the induction of endoplasmic reticulum (ER) stress (14). GAS-acquired ASN is used by the bacteria to regulate their own gene expression and rate of proliferation, constituting a novel pathogenic strategy in which GAS modulates host metabolism for its own benefit (14). Here we show that treatment of mammalian cells with the ER stressor thapsigargin (TG) or infection with wild type (WT) GAS, but not treatment with ASN, results in the production of culture supernatants that can restore biofilm biomass accumulation to a streptolysin-null GAS mutant that is otherwise attenuated in biofilm formation on mammalian cells. Furthermore, using a mouse model of human NF, we show that the streptolysin-null mutant is significantly diminished in its ability to form biofilm-like microcolonies and to spread within soft tissue when compared to the WT strain. Most importantly, pre-treatment of mice with TG restored the ability of the streptolysin-null mutant to form microcolonies and spread in the soft tissue, underscoring the essential role of streptolysin-mediated ER stress in biofilm formation during acute GAS biofilm-associated infections.

## Results

### GAS forms biofilms on mammalian cells

To examine the potential host-pathogen interactions involved in GAS biofilm establishment during acute infection, we infected mouse embryonic fibroblasts (MEFs) with GAS at a multiplicity of infection (MOI) of 10, using the clinical strain JS95 (M14 serotype) (15). We monitored GAS biofilm biomass accumulation over time using a modified crystal violet (CV) staining protocol that distinguishes bacteria from mammalian cells (16) (Fig. 1A), and by enumerating colony forming units (CFU) (Fig. 1B). Both assays revealed a significant increase in GAS biomass accumulation from 24 to 72 hours post infection (hpi) and a steady decrease thereafter (Figs. 1A, B). To ascertain whether GAS JS95 biomass accumulation represents a common trait during GAS-host interaction, we tested mouse embryonic fibroblasts (MEFs), human foreskin fibroblasts (HFFs) and human keratinocytes (HaCaTs) as well as GAS strains of other M-types for their ability to form cell-associated biofilms. Both HFFs and HaCaTs supported GAS biomass formation albeit with different kinetics than observed for MEFs (Fig. 1C). Furthermore, both GAS MGAS5005 and JRS4 strains, of M1T1 and M6 serotypes respectively, were able to form biofilms on MEFs (Fig. 1D). Thus, the ability of GAS strain JS95 to accumulate biomass on top of mammalian cells may represent a general phenomenon in GAS-host interaction, although the biofilm formation kinetics is different among strains.

**Fig. 1.**
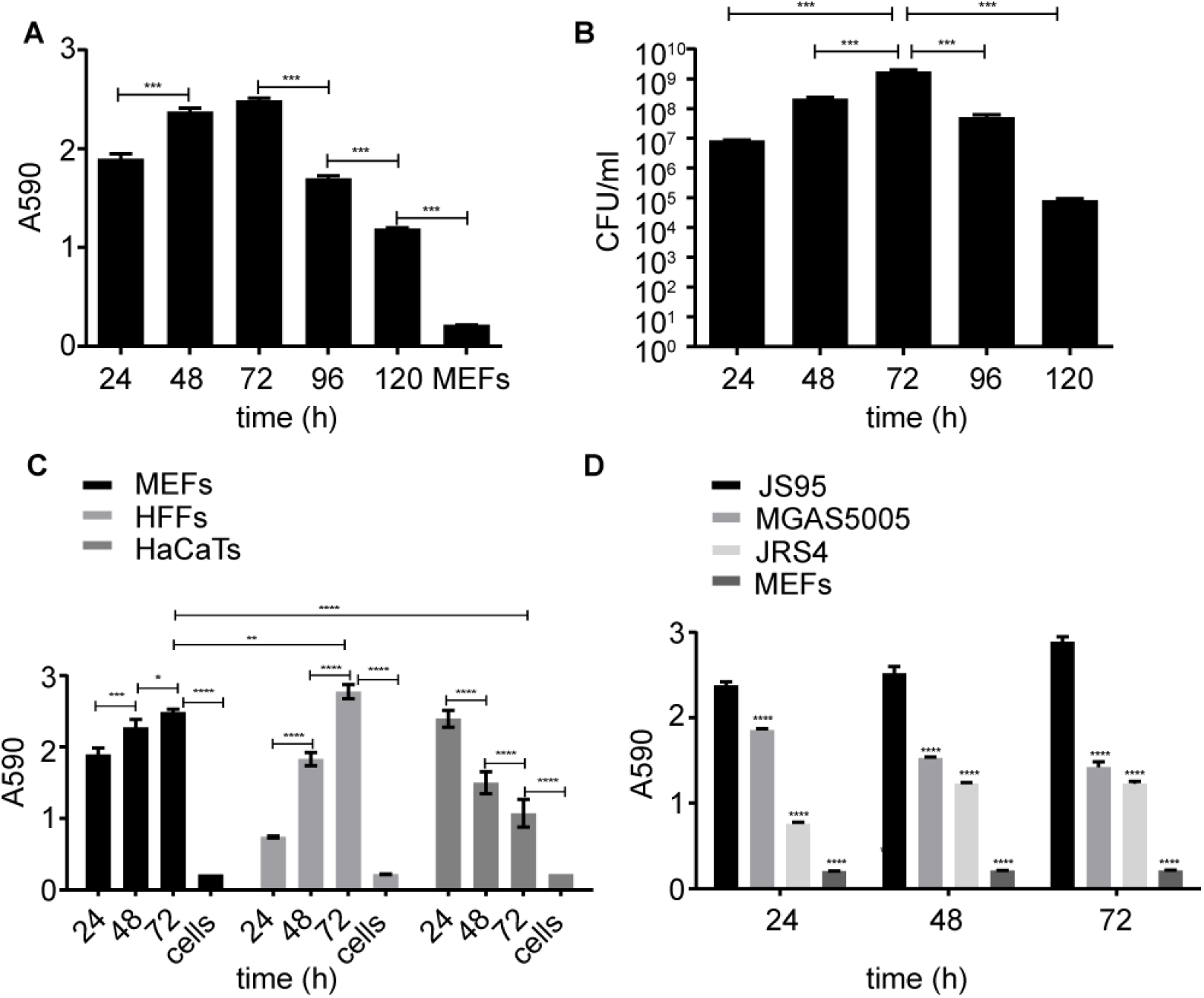
GAS forms biofilms on mammalian cells. Time course of biofilm development by GAS JS95 on mouse embryonic fibroblasts (MEFs) infected at MOI 10 and quantified by *(A)* crystal violet (CV) staining or *(B)* colony forming unit (CFU) enumeration at indicated timepoints (hours). *(C)* Quantification of biomass of GAS JS95 biofilms grown on MEFs, human foreskin fibroblasts (HFFs), and human keratinocytes (HaCaTs) by CV assay. *(D)* Biofilm biomass of GAS strains JS95, MGAS5005, and JRS4 on MEFs as quantified by CV assay. Controls represent uninfected mammalian cells. Data represent the mean ± standard error of the mean (SEM) values; N=3. Statistical significance was calculated by one-way Anova *(A, B)* or two-way Anova (C, D) using Tukey's multiple comparisons test or the uncorrected Fisher’s LSD test, respectively.

To examine the biofilm properties of mammalian cell-associated GAS, we first immunostained GAS infected MEFs with antibodies specific to GAS and imaged them by confocal laser scanning microscopy (CLSM) to study their spatial and temporal organizational dynamics (Fig. 2A). GAS biofilm biomass on MEFs increased progressively between 24 and 72 hpi and decreased thereafter (Fig. 2A), consistent with the biomass levels detected by CV staining and CFU enumeration (Figs. 1A, B). Moreover, GAS biomass growing on MEFs at 72 hpi displayed a 3-D architecture indicative of mature biofilms (Figs. 2A, **S1C, E**) (17), similar to GAS microcolony structures observed in pustule skin lesions from impetigo patients and in a mouse model of superficial human GAS skin infections (6). In addition, scanning electron microscopy (SEM) revealed that GAS biofilms on MEFs appear in chains within thick and multi-layered aggregates (Fig. 2B), similar to previous reports of GAS biofilms *in vivo* (8, 12). While JS95 is capable of forming biofilm on polystyrene tissue culture plates (referred hereafter as plastic), biofilms on this abiotic substratum accumulate less biomass compared to biofilms on MEFs, display different developmental kinetics, and have a 3-D architecture distinct from MEF-associated biofilms (Figs. S1A, *B, C, D,* E). Additionally, SEM images of abiotic-grown biofilms show reduced aggregation (Figs. S1F) compared to their MEF-grown counterparts (Fig. 2B).

**Fig. 2.**
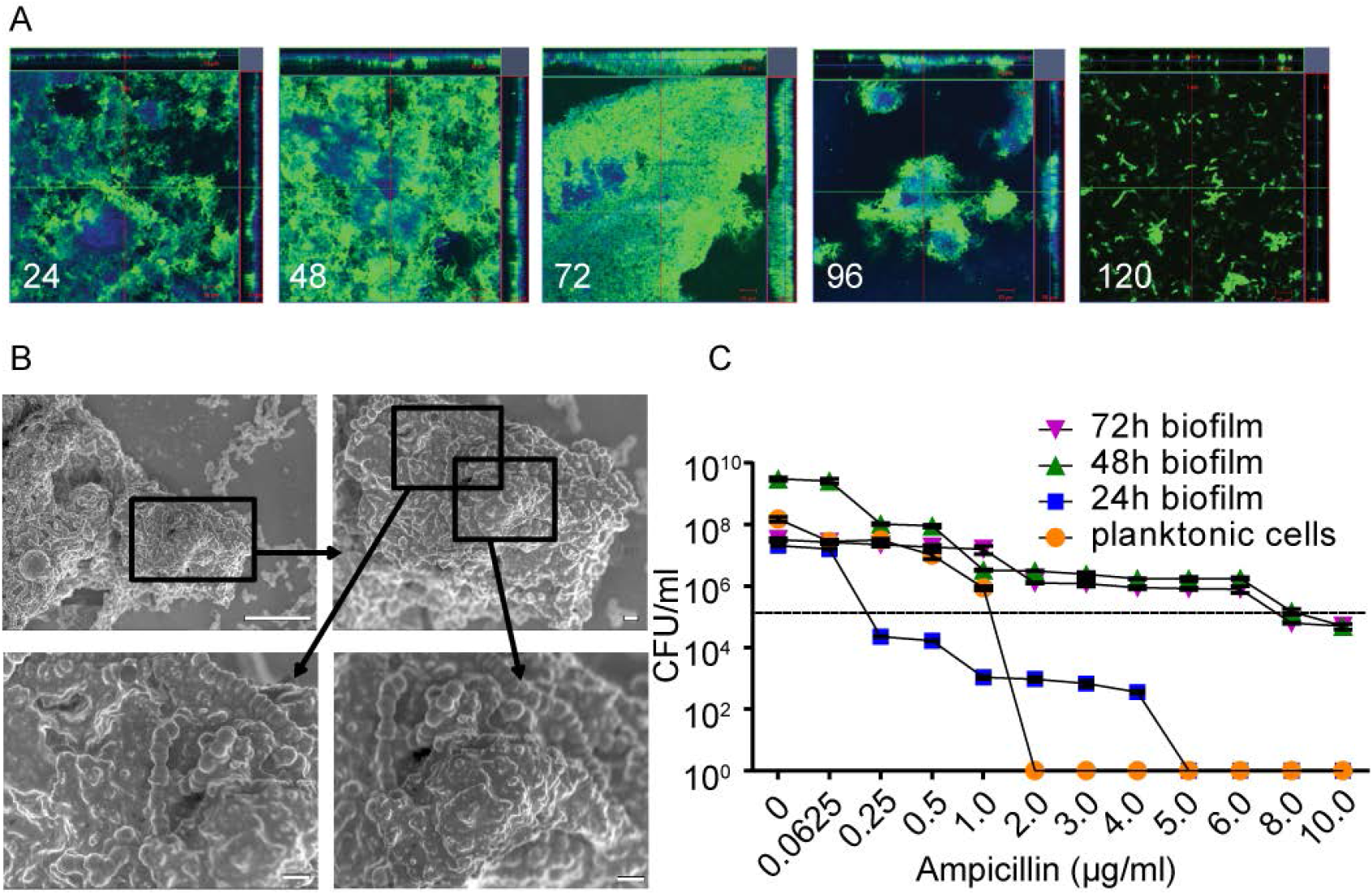
GAS growing on MEFs exhibit biofilm characteristics. (A) CLSM of GAS biofilm development and 3-D architecture on MEFs over time (hours). Bacteria were stained with anti-GAS antibody (green) and MEFs with phalloidin AF350 (blue). Orthogonal CLSM images of biofilm were acquired at x63 magnification. *(B)* SEM images of GAS biofilms grown on MEFs for 24 h at magnifications of x2500 (upper left), x5000 (upper right), and x10,000 (bottom panels). Scale bars indicate 10 µm, 1 µm, and 1 µm, respectively. *(C)* Antibiotic susceptibility of GAS in planktonic growth compared to 24 h, 48 h, and 72 h mature biofilms on MEFs after treatment with increasing concentrations of ampicillin for 24 h. Dotted line represents the initial GAS inoculum of 1 x 10^5^ CFU. Viable bacteria were quantified by CFU enumeration. Data represents the mean ± SEM values; N=3.

We next tested whether GAS growing on MEFs exhibited increased tolerance to antibiotics, a characteristic of biofilms (10, 18). We exposed pre-formed GAS biofilms growing on MEFs, or planktonic bacteria, (both seeded with an initial inoculum of 1 x 10^5^ CFU) to increasing concentrations of ampicillin and enumerated CFU after 24 h of antibiotic exposure. GAS biofilms were significantly more tolerant to ampicillin compared to planktonic cells (Fig. 2C). Taken together, these observations show that GAS forms dense host-associated biofilms during acute infection that may contribute to disease progression by enhanced tolerance towards antibiotic treatment.

### GAS streptolysin expression is necessary for biofilm formation on host cells

To identify GAS determinants important for biofilm formation on host cells, we examined the ability of isogenic mutants of JS95 WT to form biofilms on MEFs. The mutants chosen are defective in the expression of GAS virulence factors (3, 19, 20), some of which have also been implicated in GAS biofilm formation (4, 21). We observed that deletion of *emm14* encoding the M14 anti-phagocytic surface protein and inactivation of the cysteine protease *speB* enhanced biofilm formation on MEFs (**Fig. S2A**). By contrast, inactivation of *hasA,* essential for hyaluronic acid capsule production, decreased biofilm formation significantly (**Fig. S2A**). In addition, a strain lacking the expression of both SLO and SLS (*Δslo,sagI¯*) was the most highly attenuated mutant for biofilm formation on MEFs (**Fig. S2A**). Because of this marked defect of the streptolysin-null strain, we focused the remainder of our study on the role of SLO and SLS in host-associated biofilm formation. We compared biofilm formation by JS95 (WT) and the streptolysin-null mutant *(Δslo,sagI¯)* over time on MEFs versus plastic. The amount of biomass produced on MEFs by WT biofilms was significantly higher than that produced by *Δslo,sagI¯* biofilms at all time points examined (Fig. 3A). By contrast, on plastic, *Δslo,sagI¯* produced significantly more biomass at 48 and 72 hpi compared to WT (Fig. 3A). To ensure that both strains exhibit similar rates of growth in their planktonic state, we also enumerated CFU of WT and *Δslo,sagI¯* in the culture supernatants (non-biofilm fraction) while growing on MEFs and observed no defect in planktonic growth (**Fig. S2B**). To gain more spatial insight into GAS biofilms on MEFs, we next examined the architecture of biofilms formed by WT, *Δslo,sagI¯,* or *Δslo,sagI¯* harboring plasmids complementing either *slo* or *sagI*. CLSM revealed that while WT GAS aggregation on MEFs was dense, *Δslo,sagI¯* displayed sparse aggregates on MEFs (Fig. 3B). However, the corresponding complemented mutants expressing either SLO or SLS developed biofilm biomass similar to that of the WT, demonstrating that either SLO or SLS are sufficient to complement *Δslo,sagl¯* for biofilm formation (Figs. 3B, C), indicating their redundant role in host-associated biofilms.

**Fig. 3.**
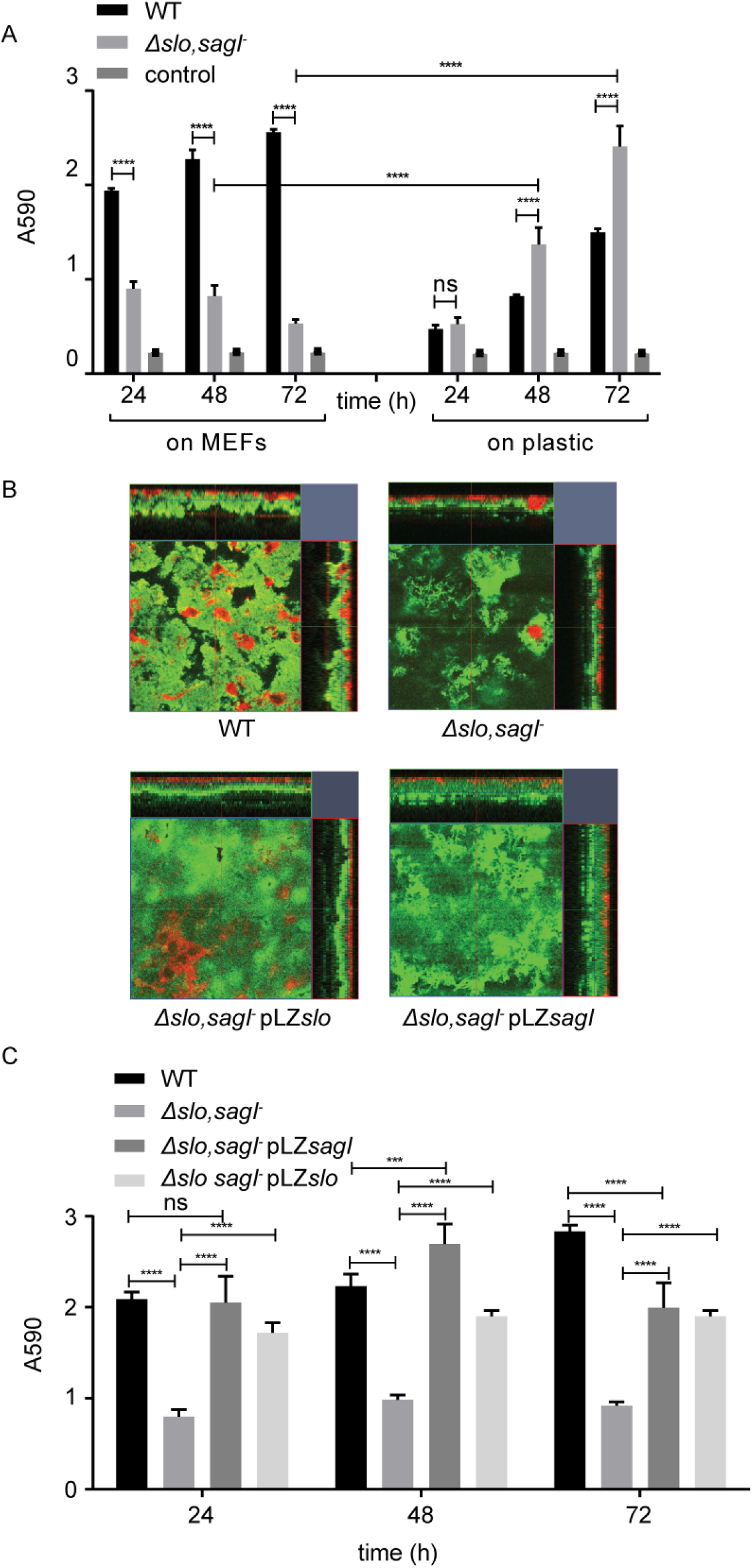
GAS streptolysins (SLS or SLO) are required for biofilm formation on mammalian cells. *(A)* Biofilm biomass accumulation of WT and *Δslo,asgI¯* on MEFs or plastic over time (hours) was quantified by CV assay. Uninfected MEFs or media alone served as negative controls. *(B)* CLSM images of 72 h biofilms formed by GAS WT, *Δslo,asgľ,* and complemented strains at x63 magnification. Bacteria are stained with anti-GAS antibody (green) and MEFs with concanavalin A AF594 (red). *(C)* Quantification of biofilm biomass formed by GAS WT, *Δslo,asgľ,* and complemented strains on MEFs was performed by CV staining. Data represent the mean ± SEM values; N=3. Statistical significance was calculated by two-way Anova using the uncorrected Fisher’s LSD test *(A, C)*.

### ER stress-induced host-associated factor(s) contributes to GAS biofilm formation on host cells

GAS streptolysins SLO and SLS cause ER stress in host cells leading to the release of asparagine (ASN) into the culture media, which GAS JS95 can sense and use to regulate its gene expression and rate of proliferation (14). Since SLO or SLS are required for biofilm formation, we hypothesized that the streptolysin-mediated ER stress pathway could similarly be involved in host cell-associated biofilm formation. To determine if ER stress induction resulted in the release of a biofilm-promoting factor, we supplemented the *Δslo,sagl¯* double mutant biofilm with filtered (0.22 uM) conditioned media (CM) from MEFs treated with the ER stressor thapsigargin (TG) for various time durations (22). We observed a significant recovery in MEF-associated biofilm formation in the presence of CM obtained from MEFs treated with TG for 36 or 48 h (Fig. 4A) whereas CM obtained from untreated MEFs did not affect biofilm formation (Fig. S3A). Prolonged or unmitigated ER stress can trigger apoptosis (23). Indeed, after 24 h of TG treatment, apoptotic MEFs started to appear and their number increased considerably after 36 h of treatment (**Fig. S4**). To examine if apoptosis of MEFs *per se* was responsible for promoting MEF-associated biofilm formation, we treated MEFs with etoposide (ET), which inhibits DNA damage repair and consequently triggers apoptosis in an ER stress-independent manner (24). We again supplemented *Δslo,sagl¯* grown on MEFs with filtered CM from MEFs treated with ET and examined biomass accumulation. In contrast to the CM of TG-treated MEFs, CM of ET-treated MEFs failed to promote biofilm formation (Fig. 4B), even after 36 h when a large population of MEFs was already apoptotic (**Fig. S4**). Neither TG nor ET affected planktonic GAS growth (**Fig. S5**). These results suggested that ER-stress leads to the release of a host-associated biofilm promoting factor or factors.

**Fig. 4.**
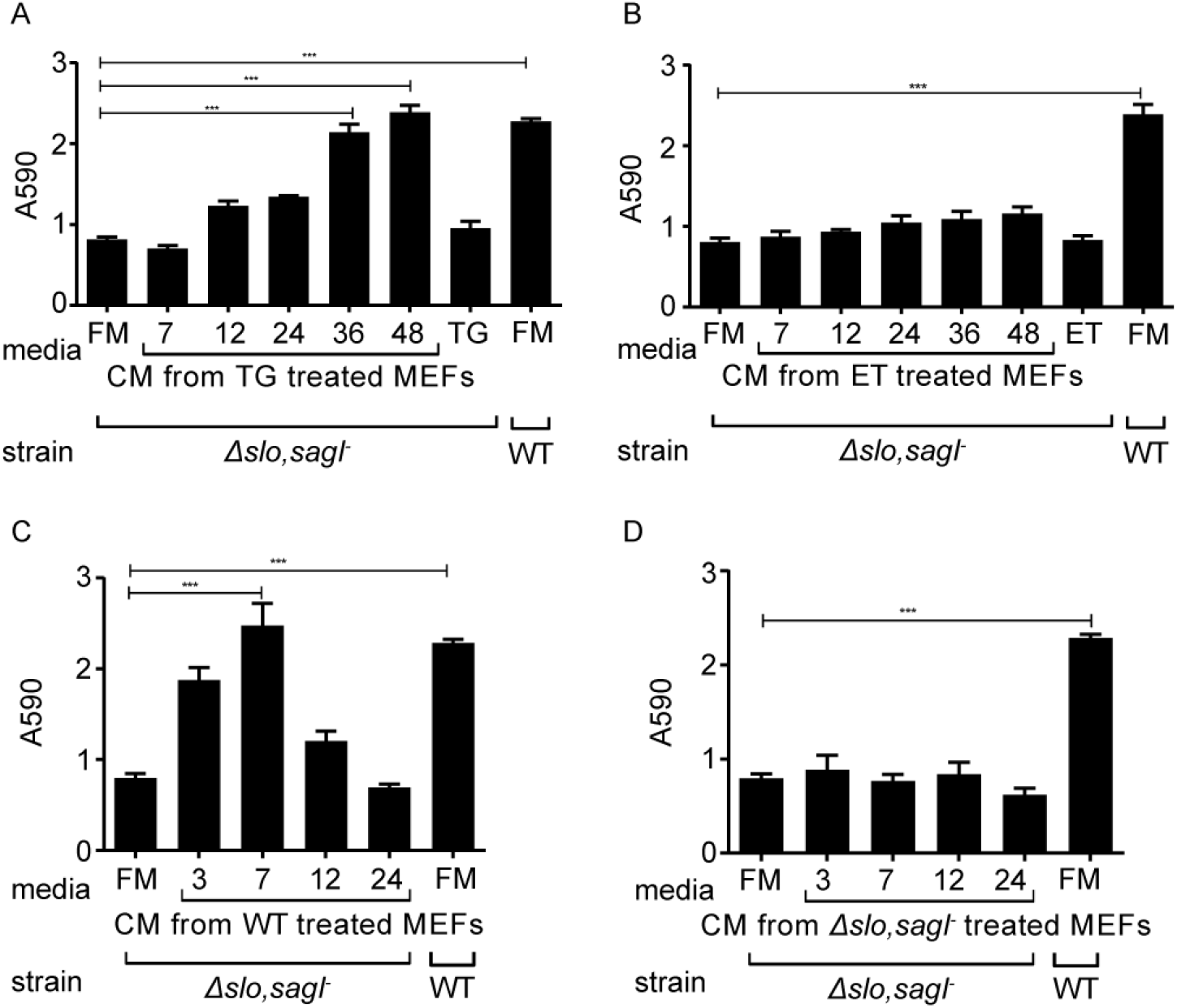
An ER-stress induced factor is required for biofilm formation on mammalian cells. Conditioned media (CM) from uninfected MEFs treated with TG *(A)* and ET *(B)* were collected at the indicated time points (numbers denote hours of treatment). Formation of biofilm by WT GAS on top of MEFs in fresh media (FM) was used as a positive control. FM alone or drug-supplemented FM (TG/ET) served as negative controls. CM from WT GAS (C) or *Δslo,asgľ* (D) on MEFs were collected at the indicated time points. *Δslo,asgľ* biofilm biomass was quantified in their presence after 24 hpi. Positive and negative controls are the same as above. Data represent the mean ± SEM values, N=3. Statistical significance was calculated by one-way Anova using Tukey’s multiple comparison test.

SLO and SLS are well characterized cytolysins that are secreted by GAS (25). Lactate dehydrogenase (LDH) release into the culture supernatants as a measure of streptolysin-mediated cytotoxicity revealed progressive cell death of WT infected MEFs starting at 4 hpi. By 10 hpi, greater than 90% cell-cytotoxicity was observed (**Fig. S6**). By contrast, the *Δslo,sagI¯* mutant caused only 30% cytotoxicity in MEFs at the same time point (**Fig. S6**). Consistent with this, we observed that WT-infected MEFs displayed markers of apoptosis as early as 6 hpi and necrosis by 8 hpi (**Fig. S7**). By contrast, neither apoptosis nor necrosis occurred in MEFs infected with *Δslo,sagI¯* during the time course examined (**Fig. S7**). Therefore, to test whether streptolysin-induced ER stress and the subsequent apoptosis cascade also elicited the release of a biofilm-promoting factor or factors, similar to TG-treated MEFs, we examined the ability of CM from MEFs infected with WT GAS to restore biofilm formation to the attenuated *Δslo,sagI¯* mutant. As a control, we used CM from MEFs infected with *Δslo,sagI¯*. CM from WT-infected MEFs collected at early times (3 hpi), yielded partial (∼80%) restoration of *Δslo,sagI¯* biofilms (Fig. 4C). By 7 hpi, when induction of the ER stress-associated gene *chop* was also observed (**Fig. S8**) (26), CM from WT-infected MEFs fully complemented *Δslo,sagI¯* biofilm formation to levels equivalent to WT (Fig. 4C). These findings suggest that a single or multiple biofilm-promoting factors are released during the GAS-mediated ER stress and apoptotic cascade. Biofilm formation was not restored by CM from MEFs infected with WT at later time points (12 or 24 hpi) (Fig. 4C), possibly due to a transient nature or activity of the released biofilm-promoting factor. CM harvested from MEFs infected with *Δslo,sagI¯* also did not restore biofilm formation to the double mutant (Fig. 4*D*) suggesting that streptolysins are essential for the release of this factor(s).

Since SLO and SLS are known to be released into the culture medium (27, 28), we anticipated that the high level of biofilm formation observed in Fig. 4C could, in part, be caused by the secreted SLO and SLS present in the filtered CM. To test this, we used culture media of WT or of *Δslo,sagI¯* grown on plastic for different time periods (**Figs. S3B, C**). As predicted, culture medium of WT GAS grown for 7 h on plastic plates partially restored MEF-associated biofilm formation of *Δslo,sagI¯,* most likely due to the presence of SLO and SLS in the medium leading to induction of ER stress in the MEFs and release of the biofilm-promoting factor (**Fig. S3B**). By contrast, culture media of *Δslo,sagI¯* grown on plastic under the same conditions failed to restore biofilm formation at any time point of growth (**Fig. S3C**). Since ASN is released by host cells upon induction of ER stress by SLO and SLS resulting in enhanced GAS proliferation (14), we tested whether ASN is involved in the restoration of MEF-associated biofilm formation. However, addition of increasing concentrations of ASN did not restore *Δslo,sagI¯* biofilm formation, indicating that under these assay conditions ASN is insufficient for complementation and that other host factors may be involved (**Fig. S3D**).

### Streptolysin-mediated ER stress promotes GAS microcolony formation and spread *in vivo*

To assess the contribution of streptolysin-mediated ER stress in GAS microcolony/biofilm formation *in vivo,* we used a murine model mimicking human GAS NF (15, 29). We injected GAS subcutaneously and assessed bacterial burden and spread across the infected soft tissue at 12 hpi. Infection with WT resulted in high titer infection of more than 1 x 10^10^ CFU/g in the soft tissue (Fig 5A). To obtain high resolution imaging of the bacteria and surrounding host tissue, we serially sectioned infected tissue biopsies, performed immunostaining, and visualized the stained sections by confocal scanning electron microscopy (CLSM) across the transverse anatomical plane. In WT-infected mice, GAS microcolonies were concentrated predominantly in the fascia (Figs. 5B and S104). By contrast, in tissues from mice infected with *Δslo,sagI¯,* we recovered significantly fewer CFU and most commonly observed only sporadic bacteria in the fascia (Figs. 5A, B). However, more extensive examination of multiple sections across the *Δslo,sagI¯* infected tissue revealed the presence of fascial microcolonies in only a few discrete regions of the tissue biopsy, while most other sections were entirely devoid of microcolonies (Figs. 5C, **S94**). This observation suggested that the streptolysin null mutant was unable to spread throughout the infected soft-tissue and instead displayed localized aggregation. To quantify the lateral spread of microcolonies across the tissue, we measured the mean fluorescence intensity (MFI) from immunostained microcolonies in evenly spaced sections across the biopsied tissue from five mice. In contrast to WT infected tissue in which GAS was evenly spread across the examined tissue, *Δslo,sagI¯* remained localized and did not spread (Figs. 5D, E; **S9A** and **S10A**).

Since CM from host cells pre-treated with TG, but not with ET, restored the ability of *Δslo,sagI¯* to form biofilms on host cells *in vitro* (Figs. 4A, B), we tested whether TG also promoted microcolony formation in soft tissue *in vivo*. Subcutaneous injection with TG administered 12 h prior to inoculation with *Δslo,sagI¯* resulted in a ∼100-fold increase in the number of viable bacteria recovered 12 hpi (Fig. 5A) and restored the ability of *Δslo,sagI¯* to form microcolonies that could now disseminate laterally across the fascia (Figs. 5C, F and **S9B**). Pre-treatment with TG did not significantly alter the lateral or transverse distribution of WT GAS microcolonies (Figs. 5G, **S10B**). However, pre-treatment with TG increased the depth of microcolony spread across the transverse plane, from the fascia into the deep muscle region for *Δslo,sagI¯* (Figs. 5C, **S9B**). Pre-treatment of mice with ET did not affect CFU recovered from the soft tissue of WT or *Δslo,sagI¯* infected mice, compared to the recovery of those strains from PBS-pretreated mice (Fig. 5A). In ET-pretreated mice, WT microcolonies and only few *Δslo,sagI¯* microcolonies were visible in the fascia and upper muscle region, respectively, and ET did not promote lateral spread of *Δslo,sagI¯* microcolonies (Fig. 5H). Moreover, the total MFI/biopsy values as a combined measure of bacterial load across the entire lateral fascial span of each tissue biopsy showed that the WT-infected tissue had significantly greater bacterial load compared to streptolysin null-infected tissue. Upon TG, but not ET, pre-treatment, we observed restoration of mutant tissue MFI values to WT levels (Fig. 5I).

**Fig. 5.**
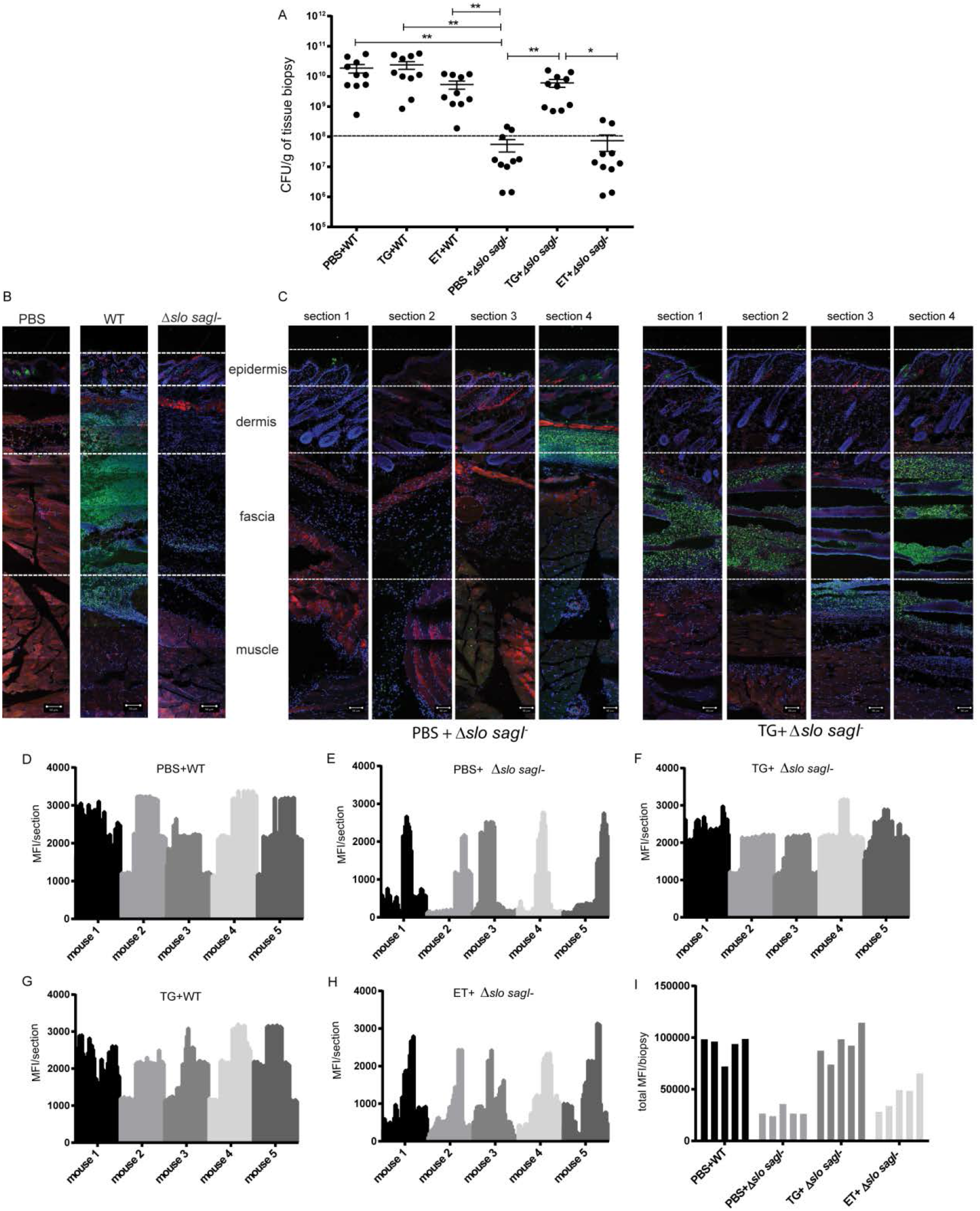
An ER-stress induced factor is required for biofilm formation on mammalian cells. GAS streptolysin-induced ER stress promotes GAS microcolony formation and spread *in vivo*. Mice were pretreated with PBS, TG, or ET for 12 h and subsequently injected with either WT or *Δslo,asgI¯*. Tissue was harvested at 12 hpi. (A) Enumeration of viable bacteria retrieved from mouse biopsied tissue from each condition. CFU values were normalized to biopsy weights. Dotted line represents initial inoculum of infection (1 x 10^8^ CFU). Data represent the mean ± SEM values; N=10. Statistical significance was calculated by two-way Anova using Sidak’s multiple comparisons test. (B) Representative images of the distribution of GAS microcolonies across the transverse plane of the infected tissue biopsy 12 hpi with PBS, WT or the *Δslo,sagI¯* mutant. GAS microcolony distribution across the transverse plane is represented by tiled x20 CLSM images of tissue sections immunostained with anti-GAS antibody (green), phalloidin AF 568 (red) and DAPI (blue); N=4. *(C)* Mice were pretreated with either PBS or TG, and were then infected with *Δslo,sagI¯*. Sections 1-4 are representative sections of GAS infected tissue biopsy from each of 4 quadrants (40 sections) from tissue from a single mouse, spanning the lateral plane of the biopsy (see Supp. Methods for detail). Scale bars represent 50 µm; N=3-4. Representative images from one mouse are shown here; images from additional animals are shown in **Fig. S9**. *(D¯H)* Quantitative analysis of GAS spread laterally across the fascia by plotting the mean fluorescence intensity (MFI) values from x20 CLSM images of anti-GAS immunostained serial sections. MFI from 40 individual serial sections (spaced 36 µm apart) per tissue biopsy spanning the lateral plane of tissue are shown; N=5. *(I)* Total mean fluorescence intensity (MFI) from 40 anti-GAS immunostained serial sections per mouse shown in *D¯F* and *H* were summed, and each bar represents total MFI per biopsy (N=5).

Finally, to confirm that both GAS- and TG-induced ER stress followed by apoptosis were associated with infection *in vivo*, we stained adjacent tissue sections for bacteria and for markers of ER stress and apoptosis. Using antibodies against the ER stress marker protein CHOP (26) or by performing TUNEL staining (30), we observed that GAS, ER stress, and apoptosis predominated in the fascia in WT infected mice Figs. 6A, B). In mice infected with *Δslo,sagI¯,* we only detected apoptotic cells in the fascial tissue, with no sign of ER stress (Fig. 6B). Pre-treatment of mice with TG, followed by injection with WT GAS intensified the ER stress staining which partially masked the few apoptotic cells present underneath (Fig. 6B). Pre-treatment with TG followed by challenge with *Δslo,sagI¯* increased the presence of ER stress markers and apoptotic cells in the fascia, together with the restoration of microcolony formation (Figs. 6A, B). By contrast, pre-treatment with ET followed by challenge with the WT or *Δslo,sagI¯* resulted in apoptotic cells in the upper layer of the muscle and the fascia, respectively, and little to no indication of ER stress, as expected (Fig. 6B). The fascia of control mice injected with TG or ET, in the absence of GAS infection, stained positively for CHOP and TUNEL or for TUNEL only, respectively (Fig. 6C). These findings show that markers of ER stress and apoptosis co-localize with sites of GAS microcolony formation. Together, these results establish that streptolysin-induced ER stress is important for GAS microcolony aggregation, distribution, and spread during the course of soft tissue infection *in vivo*.

**Fig. 6.**
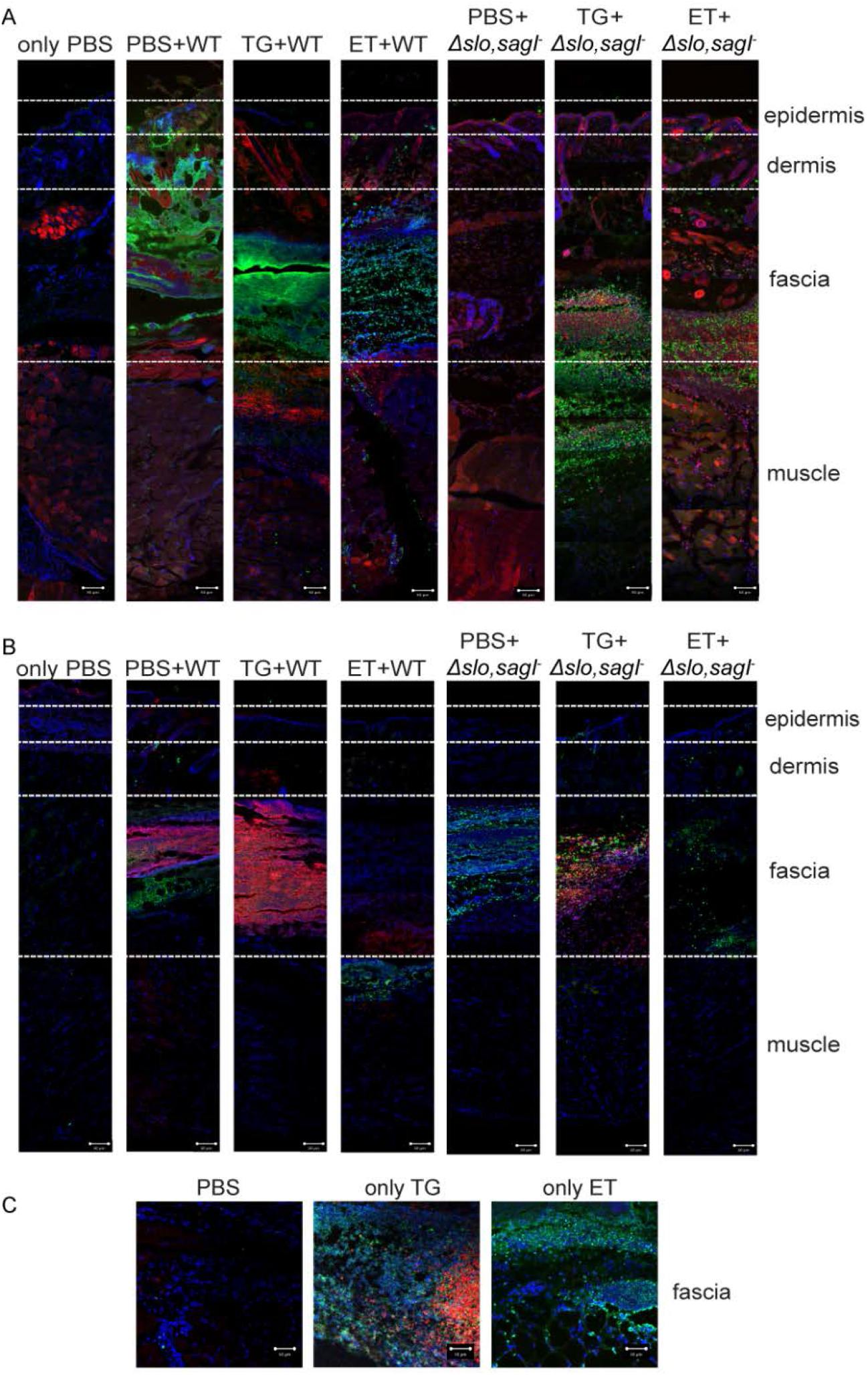
ER stress markers co-localize with GAS microcolonies *in vivo*. Mice were pretreated with PBS, TG or ET for 12 h and subsequently injected with either WT or *Δslo,asgľ*. Tissue was harvested 12 h after bacterial infection. (A) Distribution of GAS microcolonies across the transverse plane of the tissue biopsy represented by tiled x20 CLSM images of tissue sections immunostained by anti-GAS antibody (green), phalloidin AF 568 (red) and DAPI (blue). N=3-4. Representative images from one animal are shown. (B) x20 CLSM images of tissue sections adjacent to those shown in (A) after immunostaining with anti-CHOP antibody (red, for ER stress positive cells), TUNEL staining (green, for apoptosis positive cells) or DAPI (blue, all nuclei). N=2. Representative images from one mouse are shown here; images from additional animals are shown in **Fig. S10**. (C) x20 CLSM images of tissue sections (only fascia) after pre-treatment with PBS, only TG, or only ET, followed by CHOP/TUNEL co-immunostaining as described above. N=2. Representative images from one animal are shown. Scale bars represent 50 µm.

## Discussion

The contribution of biofilms to GAS pathogenesis has primarily been linked to persistence and carriage states of the bacterium (4, 21, 31). Nonetheless, the presence of dense GAS microcolonies, displaying attributes of biofilm, was identified in soft tissues of patients with acute infections such as impetigo and NF (6, 12, 15). Moreover, the presence of GAS microcolonies has also been observed in soft tissue derived from animal models of acute human GAS infections (32-34). Therefore, it is highly plausible that biofilm formation by GAS plays an important role in the development and progression of acute GAS infections such as NF. However, very little is known about the host-associated pathways and GAS determinants involved in this process.

Here, we established a model of biofilm formation on mammalian cells and showed that it recapitulates GAS microcolony formation in soft tissue during invasive GAS disease in a mouse model of human NF. Both cell-associated biofilm and host-associated microcolony formation required the expression of either SLO or SLS, and a mutant deficient for both cytolysins *(Δslo,sagI¯)* was strongly impaired in the ability to form biofilm or produce microcolonies *in vitro* or *in vivo,* respectively. In sharp contrast, GAS biofilm formation *in vitro* on an abiotic plastic surface did not require SLO and SLS, demonstrating that streptolysins are GAS factors specifically required for host-associated biofilms. This demonstrates that GAS biofilm is highly dependent on the microenvironment and that *in vitro* analyses of biofilm on abiotic surfaces may be lacking cues normally present in the host environment. Hence there is a need for using physiologically pertinent models of biofilm-associated infection. Furthermore, pre-treatment of host cells and mice with the ER stress- and apoptosis-inducing drug TG, but not the ER stress-independent inducer of apoptosis, ET, resulted in the generation of a host factor and a host microenvironment that restored the ability of *Δslo,sagI¯* to produce biofilm and microcolonies *in vitro* and *in vivo,* respectively. These observations strongly support a model in which both ER stress induction and microcolony formation are controlled by a similar host-pathogen interaction.

SLO and SLS are pore forming toxins (35, 36). Depending on their local concentrations and the type of host cells with which they interact, these toxins trigger a variety of host cellular responses ranging from ER stress to apoptosis and to necrotic cell death (25, 35, 37-41). Thus, upon infection of the host with GAS, several cellular processes occur and progress simultaneously in a time- and space-dependent manner. While it is challenging to pinpoint which of these cellular processes is responsible for biofilm and microcolony formation, our data show that ER-stress is absolutely required to initiate the process, because pre-treatment of mammalian cells with TG but not ET restored the ability of *Δslo,sagI¯* to form biofilm and microcolonies. Consistent with this, immunostaining of soft tissue sections of mice for apoptotic and ER stress markers revealed that while apoptotic markers were solely present in the fascia of mice challenged with *Δslo,sagI¯,* both ER stress and apoptotic markers were identified in the infected fascia of TG pretreated mice. Similarly, prolonged exposure of MEFs to TG and concomitant ER stress was required to generate a conditioned medium that restored the ability of a streptolysin null GAS mutant to form biofilm. Taken together, these findings suggest that events associated with the progression of host ER stress to apoptosis may be required for host-associated microcolony and biofilm formation.

Importantly, we discovered that *in vivo* triggering of ER stress not only promotes GAS microcolony formation but also enhances GAS proliferation, invasiveness into deeper tissue, and lateral spread from the injection site. TG-mediated ER stress induction significantly increased the number of viable GAS recovered from soft tissue of mice infected with the *Δslo,sagI¯* mutant. Additionally, TG pretreatment enhanced the lateral as well as the transverse spread of *Δslo,sagI¯* microcolonies from fascia to deeper muscle, compared to non-TG treated animals. TG pre-treatment did not further augment WT microcolony distribution, presumably because WT GAS-induced ER stress is already sufficient for maximal infection and spread. However, to our knowledge, this is the first report that implicates SLO and SLS with these events and demonstrates that the host ER stressed microenvironment promotes favorable conditions for GAS microcolony proliferation and dissemination.

Finally, we have shown that other host factor(s) distinct from ASN promote the formation of host cell-associated GAS biofilm. Further studies are required to determine the identity of this host-derived factor. In a recent study, Tran *et al*. reported that *Pseudomonas aeruginosa* infection and host-associated biofilm formation were dependent on an intact type III secretion apparatus (42). In that study, supernatants from epithelial cells infected with WT bacteria or from cells treated with purified GAS SLO restored biofilm formation to a type III secretion mutant, suggesting that the host-derived biofilm-promoting factor that we have identified may be a broad spectrum signal for pathogenic biofilm formation, rendering it a potentially attractive point of intervention for a variety of biofilm-associated diseases.

## Materials and Methods

### Additional details

See SI text for additional details

### Bacterial strains and growth conditions

All bacterial strains used in the study are described in Table S1 (SI). See supplementary methods for GAS growth and culture conditions.

### Cell culture and media

The cell lines used in this study were mouse embryonic fibroblasts (MEFs) (14), human foreskin fibroblasts (HFFs) (43), and HaCaT human keratinocytes (44). Cells were cultivated in Dulbecco’s modified eagle medium (DMEM+ high glucose; Gibco, New York, USA) with 10% fetal bovine serum (FBS; PAA, Pasching, Austria) at 37^o^C in 5% CO_2_ in 24-well polystyrene plastic cell culture plates (Corning, New York, USA).

### Cultivation of biofilms

Overnight cultures (16 h) of GAS strains grown in Todd Hewitt yeast extract (THY) medium (Todd-Hewitt medium (Sigma-Aldrich, Missouri, USA) supplemented with 0.2% yeast extract (Becton, Dickinson, San Jose, USA) were diluted 20-fold in a 4:1 mixture of THY media and DMEM supplemented with 10% FBS and grown until early log phase (OD_600_ = 0.2). The bacterial cells were centrifuged at 5000 g for 10 minutes, and the resulting pellet was washed twice in 1X phosphate buffered saline (PBS; Gibco, New York, USA) and then resuspended in 1X PBS to an OD._600_. of 0.8, equivalent to 1 X 10^s^ CFU/ml. Finally, 5 X 10^5^ CFU of bacteria were seeded into 24- well polystyrene plastic cell culture plates (Corning, New York, USA) containing DMEM with 10% FBS for growth on the abiotic surface. To observe GAS growth on biotic surface, MEFs seeded at a density of 5 X 10^4^ MEFs/well in 24-well plates were infected with the same bacterial inoculum, at a multiplicity of infection (MOI) of 10. The plates were incubated for static GAS growth at 37°C with 5% CO._2_. for the indicated times, with media change after every 24 h to prevent accumulation of toxic metabolic byproducts released during cellular growth.

### Biofilm quantitation

Biofilm biomass was quantified using crystal violet (CV) staining of adherent biofilms on plastic, as previously described (45) with minor modifications for biofilms growing on mammalian cells (16). The detailed protocol is described in the supplementary methods.

### Immunofluorescence and Microscopy

GAS biofilms were grown on 35 mm ibiTreat surface µ- Dishes (ibidi, Munich, Germany), either directly on the plastic surface or on MEFs seeded at 5 X 10^4^ cells/µ-Dish and immunostained with commercial antibodies, described in the supplementary methods. All confocal microscopy was performed with an LSM 780 inverted laser scanning confocal microscope (Carl Zeiss, New York, USA), equipped with a x20 Plan-Apo 0.8 NA or x63 Plan-Apo 1.4 NA oil immersion objective. Epifluorescence images were acquired using the Axio Observer Z1 Inverted Widefield microscope (Carl Zeiss, New York, USA) using a 20x/0.50 DICII and 63x/1.25 DICIII oil immersion objective. 3-D reconstruction of biofilm images and quantification were performed using the Zen Black imaging software (Carl Zeiss, New York, USA).

### Image analysis

Imaris (Bitplane AG, Belfast, United Kingdom) Version 8.4 was used to calculate bio-volume of the biofilm. ImageJ (NIH, Maryland, USA) was used to quantify mean florescence intensity of biopsy sections after immunostaining.

### Cell death assays

Cell cytotoxicity was assessed using the LDH cytotoxicity determination kit (Clonetech, California, USA) according the manufacturer’s directions. See supplementary methods for additional details. Cell death was also assessed using a NucView 488 and RedDot 2 Apoptosis and Necrosis Kit (Biotium, California, USA). Culture media was removed from infected MEFs at the indicated timepoints, cells were washed with 1X PBS, and stained according to the manufacturer’s protocol.

### Animal experiments

GAS WT or *Δslo,asgI¯* mutant strains were injected in a murine model of softtissue infection as described previously (11, 46, 47). TG or ET were injected subcutaneously into mouse soft tissue 12 h prior to the bacterial infection. See supplementary methods for additional details. All procedures were approved and performed in accordance with the Institutional Animal Care and Use Committee (IACUC) of National University of Singapore (NUS) (IACUC protocol no:R14-0575) and Nanyang Technological University (NTU), School of Biological Sciences (ARF SBS/NIE (A0294)).

### Statistical analysis

All experiments were repeated independently at least three times, with three technical replicates each (unless otherwise stated). Statistical significance for data from all experiments was determined using the GraphPad Prism software (Version 6.05 for Windows, California, United States). Data are expressed as mean ± SEM (unless stated otherwise) and p < 0.05 was considered significant (p * 0.05 (*); p < 0.01 (**); p < 0.001 (***); p < 0.0001 (****); “ns” denotes no statistical significance.

## ACKNOWLEDGEMENTS

The authors acknowledge financial support from the Singapore Centre for Environmental Life Sciences Engineering (SCELSE), whose research is supported by the National Research Foundation (NRF) Singapore, Ministry of Education (MOE), Nanyang Technological University (NTU) and National University of Singapore (NUS), under its Research Centre of Excellence Programme. This work was supported by: the National Research Foundation under its Singapore NRF Fellowship programme (NRF-NRFF2011-11); the National Medical Research Council under its Clinical Basic Research Grant (NMRC/CBRG/0086/2015), and by the National Research Foundation, Prime Minister’s Office, Singapore under its “Campus of Research Excellence and Technological Enterprise (CREATE) programme”.

We thank Ajai Vyas (NTU, Singapore) and Tan Nguan Soon Andrew (NTU, Singapore) for the kind gifts of human foreskin fibroblasts and human keratinocytes, respectively. We also thank Catherine Chen Youting (CREATE, NUS, Singapore) for technical assistance and discussions, as well as Goh Hwee Mian Sharon (SCELSE, NTU, Singapore) for assistance with *in vitro* biofilm assays. We thank Baruch B. Herzog, Miriam Ravins and Yael Kaufman (HUJ, Israel), and Artur Matysik and Ho Foo Kiong (SCELSE, NTU, Singapore) for their critical review of this paper.

